# The anaerobic WOR-3 Bacteria: Hydrogenotrophic metabolism and unique carbon fixation via Archaeal form III RuBisCO

**DOI:** 10.1101/2025.06.09.658677

**Authors:** Jianxiong Zeng, Wenzhe Hu, Licao Chang, Zhengshuang Hua, Geng Wu, Jun Liu, Guowei Wang, Chunqiao Xiao, Yun Fang

## Abstract

The uncultured WOR-3 phylum is widely distributed in anaerobic environments, including hot springs, marine ecosystems, and hydrothermal vents, yet its ecological roles and metabolic capabilities remain poorly understood. In this study, we analyzed 180 medium- to high-quality metagenome-assembled genomes (MAGs), including 59 newly reconstructed from environmental samples and 121 retrieved from the Genome Taxonomy Database (GTDB). Phylogenetic analyses resolved the WOR-3 lineage into four subgroups (subgroup 1-4). Metabolic reconstruction revealed significant divergence of the carbon, sulfur, nitrogen and hydrogen metabolism pathways among the different subgroups. Subgroup 1 was characterized by fermentative metabolism involving formate and ethanol, and uniquely exhibited potential for carbon fixation via Calvin cycle, as indicated by the presence of ribulose-1,5-bisphosphate carboxylase/oxygenase (RuBisCO) gene. Notably, WOR-3 RuBisCO is phylogenetically affiliated with archaeal form III, although the carbon fixation pathway follows the canonical bacterial Calvin cycle—a feature of potential evolutionary significance. Subgroup 3 exhibits metabolic versatility, including genes for dissimilatory sulfate reduction, sulfur oxidation, partial denitrification, and fatty acid degradation. In addition, all subgroups harbored key components of hydrogen metabolism, including widespread NiFe hydrogenases and Rnf complexes, supporting H₂-dependent electron transfer and energy conservation under anaerobic conditions. Featuring near-universal presence of complexes I and II and frequent occurrence of terminal oxidases (*cyd*AB and *cox*ABC)—suggests a facultative anaerobic lifestyle. Collectively, this study expands the genomic framework for the WOR-3 phylum and provides novel insights into the metabolic versatility and ecological functions of this previously uncharacterized lineage in biogeochemical cycles of carbon, nitrogen, and sulfur.

**IMPORTANCE:** The WOR-3 phylum represents a widespread but poorly understood bacterial lineage inhabiting diverse anaerobic/microaerobic environments. By integrating 180 metagenome-assembled genomes, including 59 newly reconstructed, this study provides the most comprehensive genomic framework to date for WOR-3. Phylogenomic and metabolic analyses revealed four distinct subgroups with divergent capacities for carbon, sulfur, and nitrogen metabolism. Notably, subgroup 1 encodes a complete Calvin-Benson-Bassham (CBB) cycle featuring an archaeal-type form III RuBisCO, suggesting an unusual evolutionary trajectory for carbon fixation in this lineage. Subgroup 3 exhibits versatile metabolic potential, including dissimilatory sulfur metabolism, partial denitrification, and fatty acid degradation, highlighting its possible roles in multiple biogeochemical processes. The widespread presence of hydrogenases and respiratory complexes across all subgroups supports energy conservation under anaerobic or microaerobic conditions. These findings not only expand the taxonomic and functional landscape of the WOR-3 phylum but also offer key insights into its ecological roles in global element cycling.

## Introduction

The capacity to culture a negligible proportion of microorganisms in their natural environment is achievable, but there is still a substantial number of uncultured microorganisms yet to be identified. The bacterial phylum WOR-3, initially discovered in hot spring environments, exemplifies this uncultured microorganism. As research advanced, the ecological adaptations of this phylum garnered increased attention. The earliest report of WOR-3 dates back to 1994, when researchers obtained the 16S rRNA sequence of WOR-3 from Octopus Spring in Yellowstone National Park, USA, and classified its species origin as ‘Candidatus Hydrothermota’ (1). In order to explore the role of WOR-3 in biogeochemical cycles, researchers have investigated its metabolic capacity. The advent of high-throughput sequencing technology has rendered metagenomic analysis an indispensable instrument in the analysis of uncultured microorganisms. A 2016 publication in the field of estuarine sediments reported that the genome of the metagenome spliced WOR-3 exhibited metabolic potential for the degradation of fatty acids and hydrogen utilization. The authors of the study suggested that this organism should be designated ‘Candidatus Stahlbacteria’ (2). However, the assembled WOR-3 genome in this literature was less than 60% complete, resulting in a failure to fully reveal its metabolic potential. Subsequently, in an anaerobic reactor environment, WOR-3 was reported to be capable of degrading amino acids (serine and alanine) and generating hydrogen (3). Furthermore, the fermentation functions of WOR-3 have been demonstrated in a simulated methane-producing and sulfate-reducing enrichment from the Jilin oilfield in China, in an anoxic environment, and three WOR-3 MAGs isolated from this environment have been shown to possess specific metabolic capabilities. It is noteworthy that the strains under scrutiny are capable of both glycolysis and gluconeogenesis pathways, as well as pyruvate oxidation and acetyl coenzyme A synthesis, in addition to ethanol fermentation and acetic acid production. It is further noteworthy that strain WOR-3 metabolizes hydrogen in the presence of [FeFe] hydrogenases of group A. However, it should be noted that WOR-3 lacks the complete tricarboxylic acid cycle (TCA cycle) and the β-oxidation pathway, which suggests that it is incapable of complete organic acid metabolism or fatty acid degradation (4). In the context of methanogenic and sulphate-reducing microbial communities, WOR-3 provides the necessary reducing power for other microorganisms (*Thermodesulfovibrio spp*.) by converting intermediate metabolites (acetic acid) into carbon dioxide and hydrogen gas (5). The ability of WOR-3 to utilize small-molecule organic matter (ethanol and acetic acid) and to rely on H_2_ metabolism for survival is well established. Nonetheless, the capacity of WOR-3 to synthesize fatty acids remains highly controversial. Given the very limited number of high-quality genomes of WOR-3 that are currently available, its ecological functional diversity, habitat distribution and environmental adaptations remain to be explored. To address these issues, we retrieved fifty-seven genomes from anaerobic sediments (56 from hot spring and 1 from lake), two genomes from an anaerobic bioreactor and 121 genomes from published genomes. By integrating existing databases with newly sampled and assembled WOR-3 genome data, this work presents an integrated and comprehensive analysis of the phylogeny, global habitat distribution, and metabolic potential of WOR-3.

## Materials and methods

### Sampling, DNA extraction, and sequencing

The sampling expedition was conducted from Jan. 2021 to Aug. 2023. Surface sediment samples were collected from the Tengchong and Quzhuomu hot springs, China. These sediments were collected into 50 ml sterile centrifuge tubes and immediately kept on dry ice until they reached the laboratory. Genomic DNA was extracted from each 10 g sediment sample using a modified phenol-chloroform method (6). A standard shotgun library with an insert size of 300 bp was inserted and then sequenced on the Illumina novaseq platform (paired-end 150-bp mode).

### Metagenomic analysis

Raw reads were pretreated using a custom Perl script and Sickle as previously reported (7). Then the resultant high-quality reads for each sample were assembled independently using SPAdes (version 3.15.2). The scaffolds were binned based on the tetranucleotide frequencies and scaffold coverage using MetaBAT (version 2.12.1) (8) with the parameters “-m 2000 --unbinned”. The preliminary classification of all bins was confirmed using the Genome Taxonomy Database Toolkit (GTDB-Tk) (9), and genome bins belonging to the WOR-3 phylum were selected. The retrieved genomes were de-duplicated using dRep software (10) and low-quality MAGs were removed, resulting in 180 medium-to-high-quality MAGs (59 genomes from our samples and 121 genomes from public database). As described previously (7), they were re-assembled using the recruited reads through BBMap (11) and were examined manually to remove possible contamination. Their completeness, contamination, and strain heterogeneity were evaluated by using CheckM (12). These curated genomes were used for the subsequent analyses including functional annotation, phylogenomic and phylogenetic analyses, and metabolic inference. The average amino acid identity (AAI) and average nucleotide identity (OrthoANI) of these 180 MAGs were then calculated using EzAAI (13) and OrthoANI (14), respectively.

### Genome annotation and metabolic reconstruction

Gene prediction of MAGs was performed using Prodigal (V2.6.3) software in ‘-p single’ mode (15). The protein-coding genes were annotated based on comparisons with the NCBI-nr, KEGG (16), EggNOG (17) and Pfam databases using DIAMOND with E-value ≤ 1e^−5^ (18). In consideration of the annotation results outlined above, a systematic reconstruction of the metabolic pathways of each genome was conducted, with a particular emphasis on pivotal metabolic pathways such as carbon metabolism (including carbon fixation), sulfur metabolism, and nitrogen metabolism.

### Phylogenomic and phylogenetic analysis

Phylogenomic tree was constructed based on a concatenation of 120 bacterial marker proteins using IQ-TREE with the parameters “LG+F+L+L+R9 -alrt 1000 -bb 1000” (19). Moreover, 16S rRNA gene sequences from WOR-3 genomes and environmental 16S rRNA gene sequences were aligned using SINA (20), and then the alignment was filtered by TrimAL with parameters set to “-gt 0.95 -cons 50” (21). 16S rRNA gene-based phylogenetic tree was constructed using IQ-TREE with the parameters “SYM+I+I+R6”. Furthermore, phylogenetic tree of *rbc*L subunit sequences was constructed using IQ-TREE with the parameters “LG+R6” and “LG+I+I+R6”, respectively. The constructed phylogenetic trees were then visualized using iTOL to display and interpret the analysis results more clearly (22).

## Results and discussion

### Genomic diversity and biogeography of WOR-3

To decipher the physiology of WOR-3, the 59 MAGs retrieved in this study and another 121 MAGs published previously were analyzed (**Table S1**). The lengths of these genomes ranged from 0.83 Mb to 4.19 Mb, the completeness ranged from 50.59% to 97.8%, and the contamination levels were observed to be less than 7.33%. Phylogenomic analysis revealed that the 180 genomes were classified into four subgroups, which was supported by the GTDB analysis and ANI/AAI analyses (**Fig. 1, 2**). The number of each subgroup is as follows: subgroup 1 (n_MAGs_=55, n_samples_=29), subgroup 2 (n_MAGs_=31, n_samples_=9), subgroup 3 (n_MAGs_=46, n_samples_=18), subgroup 4 (n_MAGs_=48, n_samples_=3). In order to comprehensively understand the ecological distribution characteristics of WOR-3, this study traced the environmental origin of WOR-3 based on the 16S rRNA gene sequences (**Table S2**) and the information of WOR-3 MAGs available in the GTDB database. It is difficult to obtain the complete 16S rRNA gene because of the incompleteness of MAGs. Some species in the phylogenetic tree based on 16S rRNA gene do not match precisely with the phylogenomic tree (**Fig. S1**). The results showed that since the discovery of WOR-3 from the hot spring environment of Yellowstone Park, USA, in 1994, more than 39 samples of WOR-3 were successively discovered in different environments from 2004 to 2011 by 16S rRNA gene sequence analysis. The application of high-throughput sequencing technology has facilitated the successful identification of metagenomic sequences of WOR-3 through the assembly of DNA fragments from environmental samples. Since 2015, the assembly of the WOR-3 genome has been achieved, leading to a substantial increase in the number of MAGs of WOR-3. The habitats of WOR-3 are highly diverse, with a primary presence in aquatic environments, including hot springs (n=62), hydrothermal vents (n=36), marine environments (n=35), freshwater sources (n=17), wastewater (n=4), and hypersaline water environments (n=4). These environments are not exclusive, however, as WOR-3 has also been identified in terrestrial habitats, including estuarine sediment (estuary sediment, n=10), terrestrial mud volcanoes (terrestrial mud volcano, n=2), oil reservoirs (oil reservoir, n=2), and bioreactors (bioreactor, n=2) (**Fig. 2**. **Table S2**). Each of the four explored subgroups of WOR-3 has been found to exist in both aquatic (e.g., hydrothermal vents, hot springs, oceans, etc.) and terrestrial environments (e.g., crustal, terrestrial mud volcanoes, etc.), but the vast majority of the genomes are predominantly distributed in aquatic environments. In terms of latitudinal distribution, WOR-3 is mainly concentrated in the middle and low latitudes (41.77°S-68.35°N).

**Fig. 1.**
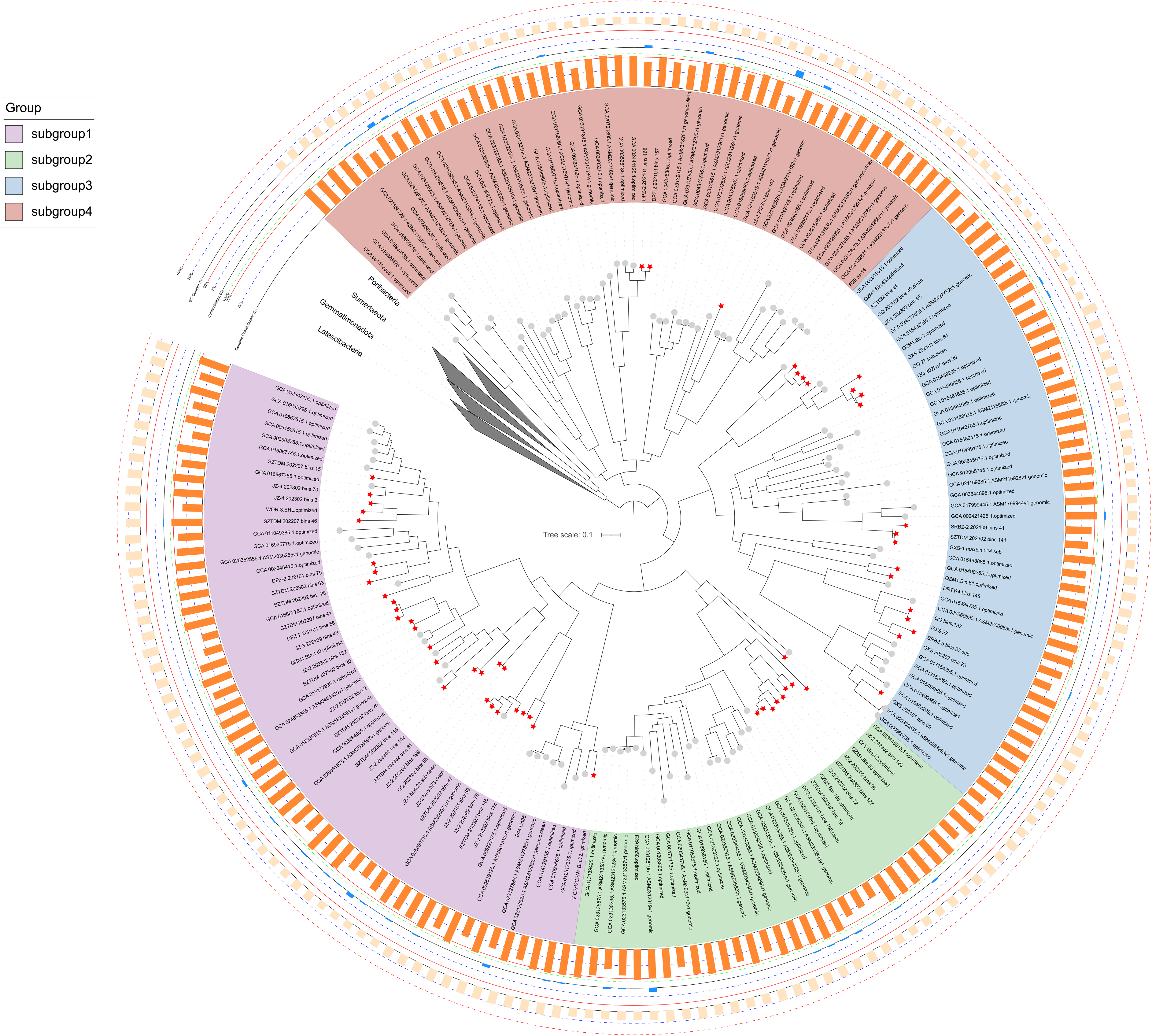
The phylogenomic tree of WOR-3 constructed based on 120 conserved marker proteins. The phylogenetic tree of the WOR-3 clade is formed by four main branches corresponding to four subgroups, and the genealogy of each subgroup is marked by different colored sectors. The distribution of sample MAGs and public database MAGs are indicated by red pentagrams and grey solid dots, respectively. Orange bars indicate the completeness of MAGs, blue bars indicate the contamination level of MAGs, and the outermost bars indicate the GC content of MAGs. Bootstrap values for phylogenetic trees greater than 75 are marked with grey solid black-edged rectangles. The genomes of *Poribacteria*, *Sumerlaeota*, *Gemmatimonadota* and *Latescibacteria* were used as outgroups.

**Fig. 2-.**
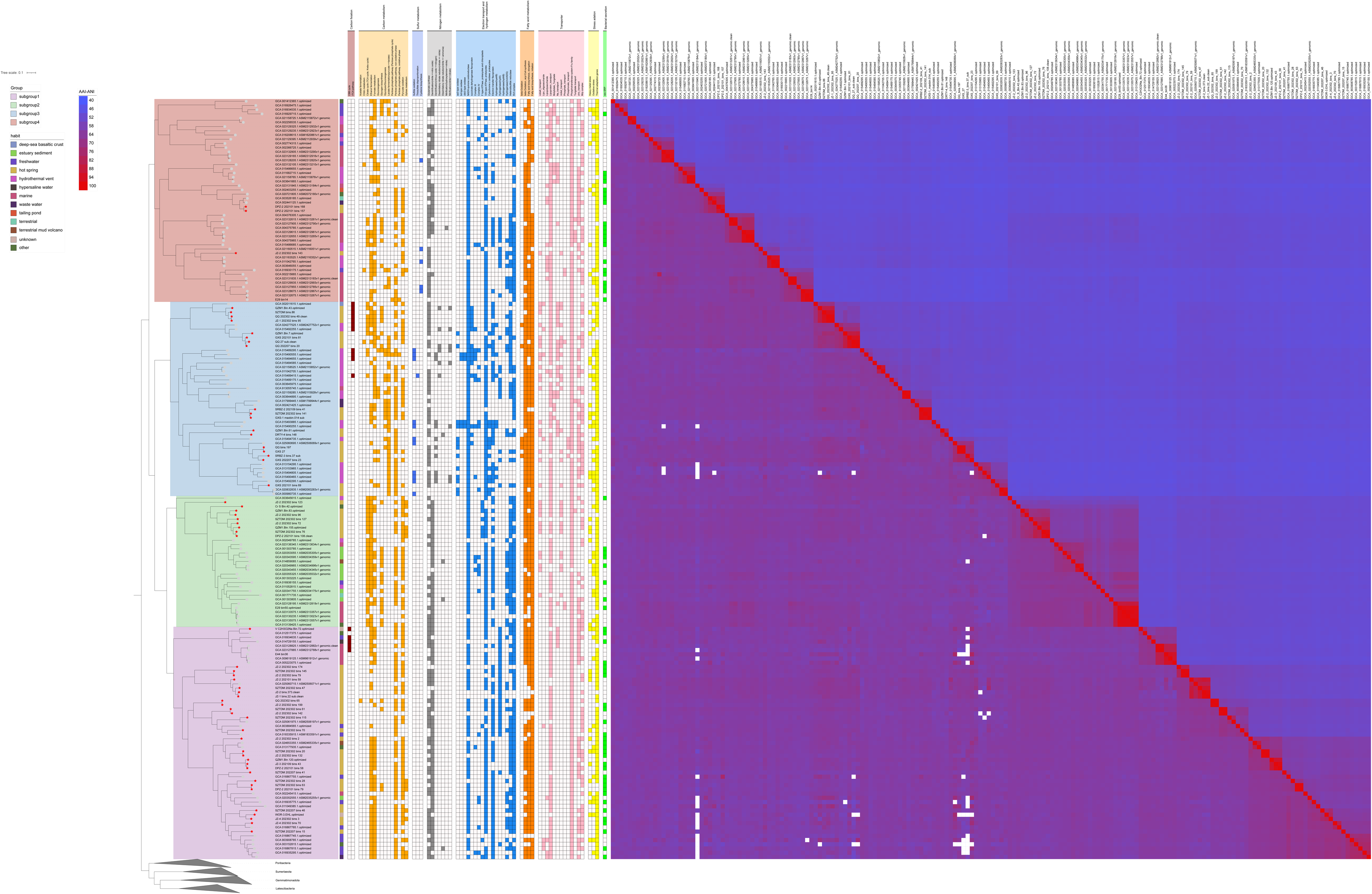
Metabolic-functional mapping, phylogenomic tree, and habitats of WOR-3. Metabolic function presence status is marked with rectangular squares of different colours and absent metabolic functions are white rectangular squares. The metabolic function mapping includes a total of nine functional categories: carbon metabolism, carbon fixation, sulfur metabolism, nitrogen metabolism, electron transport and hydrogen metabolism, fatty acid metabolism, transporter proteins, stress adaptation and bacterial secretion. The reliability of the phylogenetic tree is supported by the ANI-AAI matrix heatmap on the right, and the white squares in the heatmap indicate outliers calculated by ANI.

The extensive environmental distribution indicates the metabolic diversity of WOR-3 and its strong adaptive capacity of these little-known microorganisms to adapt to both normal and harsh environments (**Fig. 3**). Its tolerance to temperature is impressive, such as 67℃ volcanic hot springs in southwest Iceland (23) and 75-93℃ hot springs in Yellowstone National Park (24, 25). WOR-3 was similarly abundant in the hot spring (>50℃), with the relative abundance of ∼0.1% (3). The abundance of WOR-3 exhibited substantial variation across diverse reservoir environments, with significantly increased levels observed in sulfate-rich oil degradation environments (4.19-6.05%) (4), suggesting a anaerobic lifestyle and a potential role in sulfur cycling.

**Fig. 3-.**
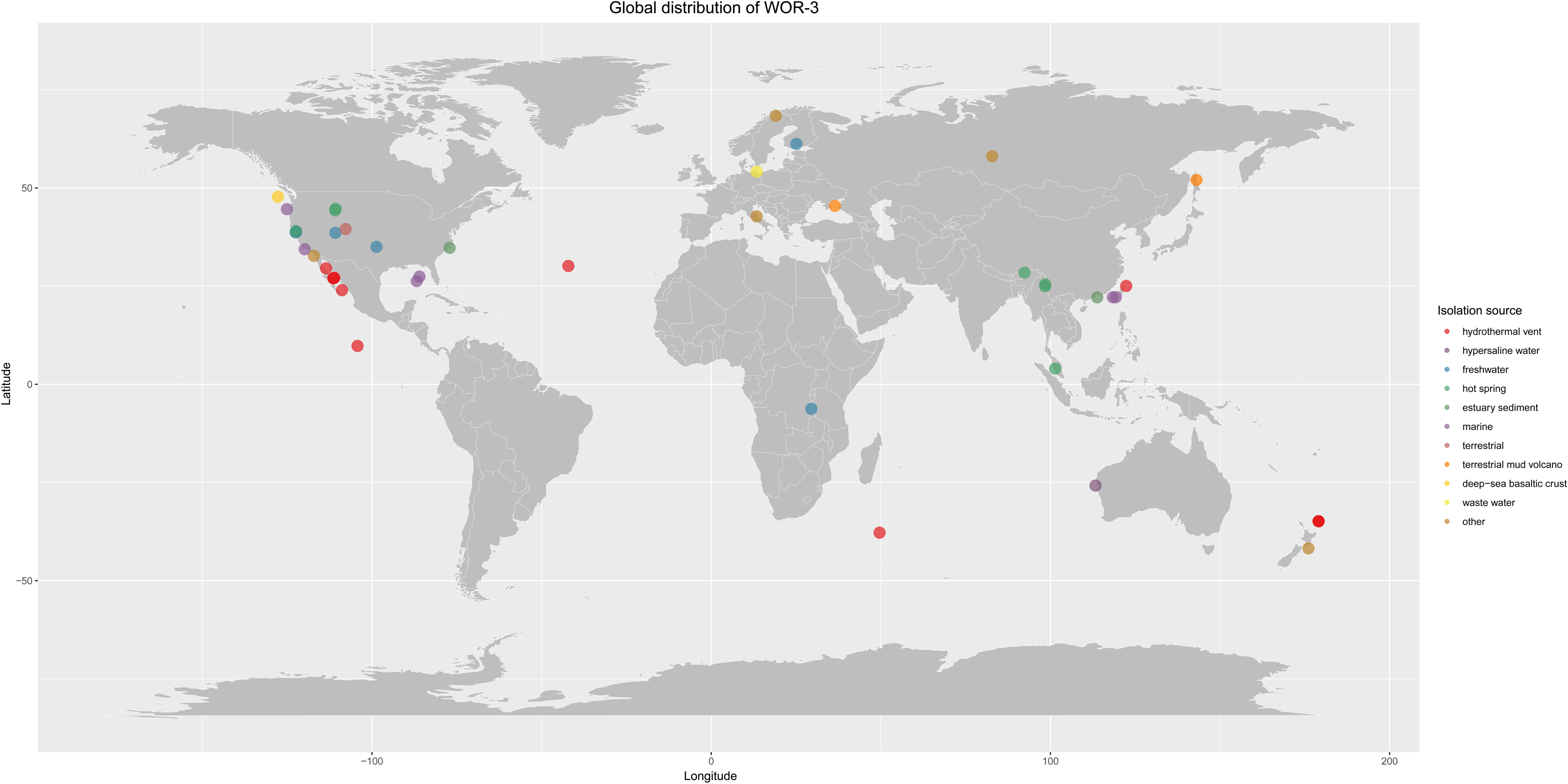
The global distribution map of WOR-3. Solid circles indicate the global distribution of MAGs, different colored circles represent different habitats of WOR-3.

### Carbon metabolism

The central carbon metabolism pathway, including glycolysis/gluconeogenesis, the pentose phosphate pathway, and the tricarboxylic acid (TCA) cycle, provide the carbon skeleton for WOR-3 and generate ATP and NADH through substrate-level phosphorylation (26). It is argued that the glycolysis/gluconeogenesis pathways and TCA cycle of WOR-3 are more complete and are capable of obtaining energy through substrate-level phosphorylation. The glycolysis pathway is subject to three rate-limiting enzymes: glucokinase (*glk*), 6-phosphofructokinase (*pfk*), and pyruvate/orthophosphate dikinase (*ppdk*) (27). The presence of dikinase (*ppd*K) is an important marker in determining the presence of this metabolic pathway. In the present study, glucokinase and pyruvate/orthophosphate dikinase were annotated in almost all genomes (**Fig. 4**. **Table S3**). However, the deletion of 6-phosphofructokinase in subgroup4 resulted in only one genome in this group with intact glycolytic capacity, a phenomenon that is similar to that observed in the thermophilic archaeon Candidatus Caldiarchaeum subterraneum (28).

**Fig. 4-.**
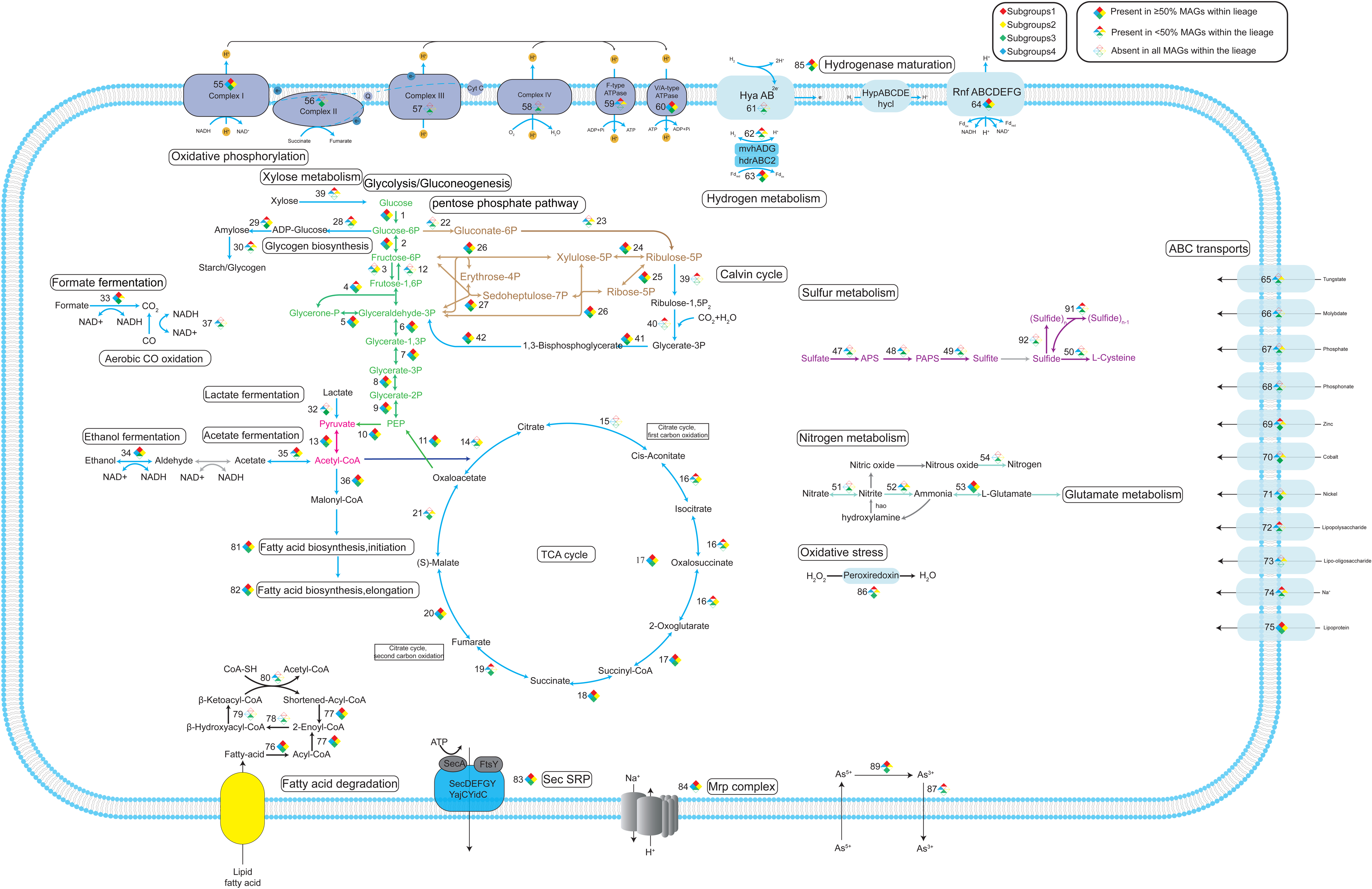
Metabolic network diagram for WOR-3. The four coloured diamonds indicate different subgroups. The prisms have three states, solid diamonds indicate that the corresponding genes are present in ≥50% of the MAGs of the subgroup, triangles indicate <50% and >0%, and hollow prisms indicate that the corresponding genes are missing from all MAGs of the subgroup. TCA: tricarboxylic acid cycle; For full names and copy numbers of the genes in number see **Table S3**.

The gluconeogenesis pathway shares a number of enzymes with the glycolysis pathway, and some of the enzymes involved in irreversible reactions (phosphoenolpyruvate carboxykinase (*pck*A), fructose-1,6-bisphosphatase (*FBP*), and glyceraldehyde 3-phosphate dehydrogenase (*GAPDH*) are key enzymes in this pathway (29). It was further demonstrated that phosphoenolpyruvate carboxykinase was classified as both GTP-dependent and ATP-dependent. The analysis revealed that these enzymes evidenced a distinct preference for different subgroups, with the GTP-dependent enzyme predominantly found in subgroup2 and the ATP-dependent enzyme predominantly distributed in subgroups1 and 3. The study concluded with the observation that only one genome in subgroup4 contained both enzymes. Despite the fact that the majority of glycolysis/gluconeogenesis-related enzymes had been annotated in WOR-3, enzymes involved in the conversion of fructose 1,6-diphosphate to fructose 1-phosphate were found in low numbers in subgroup 1. The finding that fructose 1,6-bisphosphatase was annotated in only 15 genomes across all subgroups suggests that WOR-3 does not possess a complete gluconeogenesis function (**Fig. 4**. **Table S3**).

The rate-limiting enzymes of the TCA cycle include citrate synthase (*cs*) and isocitrate dehydrogenase (*IDH*), and 2-oxoglutarate ferredoxin oxidoreductase (*kor*ABCD). Applying these three enzymes as criteria for determination, the complete TCA cycle was annotated in subgroup 2 (n=24), subgroup 3 (n=11) and subgroup 4 (n=9) (**Table S4**). This study found that combining the results of the annotation of glycolysis/gluconeogenesis and TCA cycle, the four subgroups of WOR-3 showed significant differences in the central metabolic pathways. It is clear that subgroup1 lacks the ability of glycolysis/gluconeogenesis and TCA cycle, suggesting that it is unable to harness glucose as a carbon source; whereas subgroup 2 and subgroup 4 may have the ability of intact glycolysis/gluconeogenesis and TCA cycle; and subgroup 3 only lacks the function of gluconeogenesis, suggesting that it is able to leverage glucose for metabolism.

In the absence of the glycolysis pathway, it is hypothesized that subgroup1 may rely on other carbon sources for metabolism. The presence of ethanol dehydrogenase (*adh*) in WOR-3 was identified, with the ability to oxidize alcohols to acetaldehyde, while generating the reducing power NADH and protecting cells from ethanol toxicity. This metabolic ability is prevalent in WOR-3, especially in subgroup1, where forty MAGs possess this pathway. WOR-3 has also been observed to possess the capability of converting formic acid to CO_2_ via formate dehydrogenase (*FDH*). Additionally, 33 genomes in subgroup4 have been found to contain formate dehydrogenase with coenzyme F420 as the electron acceptor. Although the small and medium subunits of carbon monoxide dehydrogenase (CO) are annotated in WOR-3, the large subunit that performs the function is not annotated, suggesting that it may not be able to fully catalyse the conversion of CO and H_2_ to CO_2_ and protons. Interestingly, 16 genomes from subgroups 1 and 2 possess xylose isomerase, which is capable of converting xylose to glucose and entering the glycolysis pathway. Among them, 11 out of 16 derived from our study, greatly expanding the number of xylose utilization members of WOR-3 (**Fig. 4**. **Table S3**). These MAGs were concentrated on the genealogical branch of the phylogenetic tree (**Fig. 2**).

Additionally, the pentose phosphate pathway (PPP) consisting of an oxidative phase and a non-oxidative phase was analyzed. The oxidative phase has been observed to be absent in many aerobic and thermophilic organisms (e.g., *Methanocaldococcus jannaschii*), whereas the non-oxidative phase has been identified as being ubiquitous in a wide range of organisms and is primarily responsible for providing RNA backbone precursors (30, 31). Key enzymes of the oxidative phase of the PPP, glucose-6-phosphate 1-dehydrogenase (*zwf*), 6-phosphogluconolactonase (*pgl*), and 6-phosphogluconate dehydrogenase (*gnd*) (27), were found in only a very small number of genomes (n=2). In addition, it was found that key genes of the non-oxidative phase were present in most genomes (n=125) (**Fig. 4**. **Table S3**). Combined with the broad distribution of WOR-3 in hot spring environments, it can be deduced that although WOR-3 is devoid of the capability to produce NADH via the oxidative phase of the PPP, it relies on transketolase (TKL) and transaldolase (TAL) as a connection to the glycolysis/gluconeogenesis pathway and continues to provide intermediary cellular metabolism such as RNA backbone metabolites.

Previous studies have reported that WOR-3 may possess the potential for fatty acid degradation(2). However, current findings indicate that this metabolic capability does not appear to be universally present in WOR-3. In WOR-3, acyl-CoA synthetase—which activates fatty acids by forming acyl-CoA during beta-oxidation—is widely distributed across all genomes. The remaining enzymes (acyl-CoA dehydrogenase, enoyl-CoA hydratase, 3-hydroxyacyl-CoA dehydrogenase, and acetyl-CoA acyltransferase) are responsible for sequentially breaking down long-chain acyl-CoA into shortened intermediates, continuously generating acetyl-CoA along with reducing power (NADH and FADH₂) (32). Notably, only six genomes within subgroup 3 encode a complete set of these enzymes required for long-chain acyl-CoA degradation, highlighting the unique metabolic role of subgroup 3 within the WOR-3 population.

### Carbon fixation

Metabolic reconstruction analyses have revealed that a proportion of the WOR-3 genome possesses carbon fixation potential, thereby challenging previous assumptions concerning the metabolic capacity of WOR-3. Specifically, five genomes from the genus B3-TA06 under subgroup 1 encode key genes in the carbon fixation pathway, including RuBisCO encoded by the rbcL/S gene, and phosphoribulose kinase encoded by the *prk* gene (**Fig. 2**. **Table S3**). It is worthy of significance that the analysis of the RuBisCO protein sequences using a phylogenetic approach demonstrated that the RuBisCO from WOR-3 is evolutionarily closer to the archaeal RuBisCO form III (**Fig. 4**). The classification of form III RuBisCO can be divided into two distinct groups: typical form III and form III B (i.e. form II/III). The typical form III is mainly from archaea, and the latter is structurally similar to the bacterial form II, but functionally similar to the archaeal form III. Phylogenetic analyses showed that RuBisCO of WOR-3 is closer to form III B (33).

Archaeal RuBisCO form III has three pathways to conduct the Calvin cycle. The first is represented by *Thermococcus kodakarensis* through the nucleoside pathway via the key enzymes AMP phosphorylase (EC2.4.2.57) and omerase (EC5.3.1.29) (33). This pathway dephosphorylates nucleotides (AMP, CMP, UMP), releasing bases to produce ribulose-1,5-bisphosphate and convertes to ribulose-1,5-bisphosphate (RuBP) by ribulose-1,5-bisphosphate isomerase (i.e., AMP → ribulose 1,5-bisphosphate → RuBP). Finally converts to 2 molecules of 3PGA from RuBP and CO_2_ via the form III or form II/III archaeal RuBisCO, participating in central carbon metabolism (34). WOR-3 unable to metabolize nucleosides via this archaeal pathway due to the absence of the key enzymes AMP phosphorylase (EC2.4.2.57) and isomerase (EC5.3.1.29). The second pathway is abiotic dephosphorylation to ribulose 1,5-bisphosphate, facilitated by archaea (*Methanocaldococcus jannaschii*, *Methanosarcina acetivorans* and *Archaeoglobus lithotrophicus*) under high temperature conditions (35). In contrast, the WOR-3 MAGs with carbon fixation potential were derived from marine, hypersaline environments, freshwater, and anaerobic bioreactors, respectively, which do not share high temperatures as a commonality. Considering that WOR-3 possesses key genes for the bacterial Calvin cycle, it is hypothesized that carbon fixation in WOR-3 still relies on the bacterial pathway. This phenomenon is not exclusive to WOR-3, but is also exhibited by *Ammonifex degensii* and *Thermodesulfitimonas autotrophica*, which belong to the phylum *Firmicutes* (36). The RuBisCO of WOR-3 is more deeply placed in the phylogenetic tree (**Fig. 5**). Besides, key genes involved in carbon fixation within the reverse tricarboxylic acid cycle (rTCA cycle) have been identified in 11 genomes from subgroup 3, encompassing citrate synthase (*gltA*) and fumarate reductase (*frd*ABCD), and 2-oxoglutarate/2-oxoacid ferredoxin oxidoreductase (*kor*ABCD) (**Fig. 4**. **Table S3**). Nevertheless, it should be noted that false-positive results cannot be ruled out on the basis of these data due to the lack of further experimental validation. It is submitted that the carbon fixation potential of WOR-3 extends the knowledge of its metabolic diversity, and moreover offers novel perspectives for understanding its adaptations in complex environments. This finding highlights the potential importance of WOR-3 in carbon fixation, and accordingly lays the foundation for further studies of its ecological functions.

**Fig. 5-.**
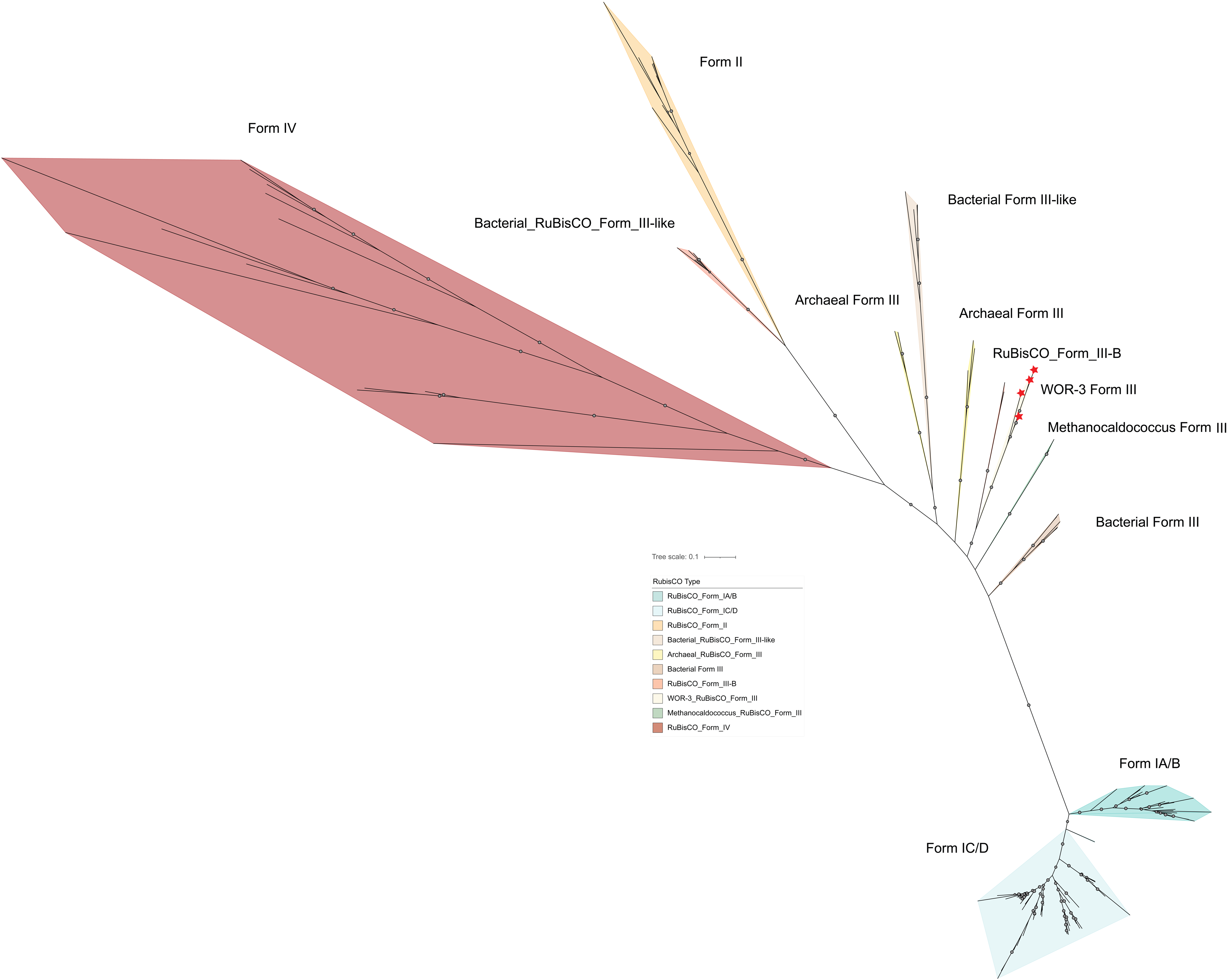
Phylogenetic tree constructed from the RuBisCO protein. RuBisCO forms 10 branches in the phylogenetic tree, and the outgroup branch is Form IV type. The branch where RuBisCO is located in WOR-3 is marked with a red pentagram.

### Oxidative phosphorylation

The energy metabolic pathways of microorganisms can be classified into two main categories: aerobic and anaerobic metabolism. Anaerobic metabolism of microorganisms is particularly important in anoxic environments such as hot springs and hydrothermal vents. In such environments, the prevalence and critical importance of nitrogen, sulfur and methane metabolism is evident, with these pathways involving a variety of electron acceptors such as nitrate, fumarate, arsenate, selenate, thiosulphate, sulfur, sulphate, oxidized metal ions and carbon dioxide (37, 38). The anaerobic metabolic processes are widespread in diverse microbial communities.

The oxidative respiratory chain consists of multiple complexes, including complex I (NADH dehydrogenase) encoded by the *nuo* gene, complex II (succinate dehydrogenase flavoprotein) encoded by *sdh*ABCD, complex III (coenzyme Q: cytochrome C oxidoreductase) encoded by *pet*B, and complex IV (cytochrome C oxidase) encoded by *cox*ABC and *cyd*AB. It was observed that most MAGs did not possess all four complexes concurrently, and only two MAGs affiliated with subgroup 3 possessed a complete respiratory chain. This finding lends further support to the hypothesis that WOR-3 is unable to transfer electrons to oxygen via the oxidative phosphorylation pathway, thereby validating its anaerobic or facultatively anaerobic lifestyle. The analysis further revealed that the majority of WOR-3 genomes encode V/A-type ATPases to produce ATP, while a comparatively smaller percentage of genomes possess oxidative enzymes, suggesting a correlation with the predominance of WOR-3 in anoxic environments.

Notably, in subgroup 3, nearly all genomes encode Complex I (*nuo* genes) and Complex II (*sdh*ABCD). Among the remaining respiratory complexes, seven MAGs contain *cyd*AB (encoding cytochrome bd ubiquinol oxidase), while 16 MAGs possess *cox*ABC (encoding aa3-type cytochrome c oxidase) (**Fig. 4**. **Table S3**). Based on these genomic features, we propose that subgroup 3 likely consists of facultative anaerobes rather than obligate anaerobes.

### Sulfur metabolism

To further explore the sulfur metabolism potential of WOR-3, we analyzed its electron receptor utilization capacity in relation to sulfur metabolism. Sulfhydrogenase (*hyd*ABDG), an enzyme that can transfer polysulfide_n_ and H_2_ into polysulfide_(n-1)_, H_2_S, and NADH, was found in 42 WOR-3 MAGs (39). The corresponding functions are attributed to subgroup 1, 3, and 4. Furthermore, the results show that 11 MAGs in subgroup 3 support assimilatory sulfate reduction (ASR), with the vast majority (n=10) coming from hydrothermal vent environments and only one from a hot spring environment. Key enzymes involved in ASR include sulfate adenylyltransferase (*sat*), adenylylsulfate kinase (*cys*C), and phosphoadenosine phosphosulfate reductase (*cys*H) (40). The enzymes are distributed in other subgroups, but it is only MAGs from subgroup 3 that possess all three enzymes simultaneously and are able to reduce sulfate to sulfite. However, WOR-3 is unable to further reduce sulfite to sulfide because it lacks sulfite reductase (*cys*JI/*sir*). With regard to the fate of sulfide, it was observed that some MAGs (n=39) possessed cysteine synthase (*cys*K) in various subgroups. These genomes lacked ASR capability, and thus could only synthesize cysteine using exogenous sulfide (**Fig. 4**. **Table S3**). Conclusively, WOR-3 could transfer electrons via sulfur metabolism through ASR pathway and sulfur hydrolases that can provide some reducing power.

### Nitrogen metabolism

The nitrogen metabolism in subgroup 3 of WOR-3 is also impressive. Dissimilatory nitrate reduction (DNR), which reduces nitrate to nitrite by nitrate reductase (*nar*GHI), was found in six genomes of subgroup 3. Subgroup 3 also exhibited 10 MAGs containing *nos*Z genes, which encode nitrous-oxide reductase (NOR), suggesting its capacity for denitrification through the reduction of nitric oxide (NO) to nitrogen gas (N_2_). WOR-3 genomes with above-mentioned nitrogen metabolism capacity mainly from subgroup 3, while not for the other subgroups (**Fig. 4**, **Table S3**). These observations suggest that subgroup 3 possesses a distinct metabolic profile within the WOR-3 clade, which may be instrumental in fulfilling its ecological function. This disparity in metabolic capacity underscores the diversification among the different subgroups of WOR-3 with respect to ecological adaptation and functional diversity.

### Hydrogen metabolism

In the MAGs of WOR-3, a variety of hydrogenase genes were identified, thus indicating that WOR-3 possesses considerable potential for H₂ metabolism. [NiFe] methyl-viologen-reducing hydrogenase, heterodisulfide reductase, and rnf complex are the most significant due to their prevalence within the WOR-3 metagenome. Specifically, 34 MAGs from various subgroups of WOR-3 possessed both [NiFe] methyl-viologen-reducing hydrogenase (*mvh*ADG) and heterodisulfide reductase (*hdr*ABC), forming a functional complex capable of simultaneous reduction of Fe-oxo-protein and CoB-CoM heterodisulfide during H_2_ oxidation. It is important to note that analogous metabolic mechanisms have been reported in hydrogenotrophic methanogens (41), as well as in certain bacteria such as *Deltaproteobacteria* and *Sumerlaeota* (42, 43). MvhADG-hdrABC2 complex in WOR-3 may be analogous to that in *Sumerlaeota* and involved in energy-efficient metabolism in the population. Similarly, the Rnf complex has been found to be widely distributed across subgroups (135/180 MAGs). The Rnf complex is a respiratory enzyme widely present in bacteria, and exists as the sole respiratory enzyme in some anaerobic acetate-producing bacteria, where it plays a crucial role in bacterial energy metabolism (44). Phylogenetic analysis of the *rnf* C subunit (**Fig. S2**) revealed its clustering with anaerobic organisms, thereby further substantiating the anaerobic lifestyle of WOR-3. These findings imply that WOR-3 may have adapted to its hypoxic survival environment by facilitating energy metabolism and electron transfer in anaerobic environments through the mvhADG-hdrABC2 complex and rnf complex. This metabolic strategy not only reveals the functional potential of WOR-3 in anaerobic ecosystems, but also provides important clues for understanding its ecological adaptations.

The four subgroups of WOR-3 share certain metabolic characteristics, including 1) an anaerobic or facultatively anaerobic lifestyle; 2) the potential for H_2_ metabolism; and 3) carbon metabolism via fermentation and non-oxidative phase of the pentose phosphate pathway. The phylum WOR-3 exhibits substantial diversity in carbon metabolism and ecological functional specialization. Comparative genomic analysis reveals clear distinctions among its four subgroups in core metabolic pathways. Subgroup 1 lacks the genetic potential for glycolysis, gluconeogenesis, and the TCA cycle, indicating an inability to utilize glucose as a primary carbon source. However, it can metabolize alternative substrates via auxiliary pathways such as the xylose isomerase route. In contrast, subgroups 2 and 4 possess complete gene sets for glycolysis, gluconeogenesis, and the TCA cycle, supporting efficient glucose utilization. Subgroup 3 encodes the glycolytic and TCA cycle pathways but lacks gluconeogenic capacity. Notably, this subgroup displays distinct capabilities in sulfur and nitrogen metabolism, including genes associated with assimilatory sulfate reduction (ASR), dissimilatory nitrate reduction (DNR), and denitrification. Additionally, two genomes within Subgroup 3 harbor a complete oxidative phosphorylation chain, potentially representing the only members within WOR-3 capable of ATP production via aerobic respiration. Carbon fixation potential is restricted to Subgroups 1 and 3, mediated through the CBB cycle and the rTCA cycle, respectively, reflecting divergent ecological strategies across the lineage.

## Conclusion

This study provides a comprehensive investigation of the metabolic potential and ecological functions of the phylum WOR-3. Phylogenetic analyses based on 16S rRNA genes and metagenome-assembled genomes (MAGs) revealed the global distribution of WOR-3 and its presence across diverse habitats, particularly in anoxic aquatic environments such as hot springs, hydrothermal vents, marine, and freshwater systems. Metabolic reconstruction of 180 medium- to high-quality MAGs showed that, while the overall metabolic repertoire of WOR-3 is relatively conserved, two major lineages—subgroup 1 and subgroup 3—exhibit marked metabolic differentiation. Subgroup 3 displays unique capabilities in sulfur and nitrogen metabolism, including dissimilatory sulfate reduction and denitrification, whereas subgroup 1 appears to rely on the assimilation of complex carbon sources (e.g., xylose) to meet its nutritional demands. Notably, carbon fixation via the CBB cycle was identified in subgroup 1 for the first time. Phylogenetic analysis further revealed that WOR-3 encodes a form III archaeal-like RuBisCO, yet its carbon fixation pathway is functionally aligned with the bacterial CBB cycle rather than the archaeal nucleotide-based pathway. This finding provides new insights into the evolution of autotrophic carbon fixation pathways.

The metabolic diversity of WOR-3 reflects distinct ecological adaptation strategies. Subgroup 1 lacks the genetic potential for glycolysis and the tricarboxylic acid (TCA) cycle, yet can metabolize alternative carbon sources through auxiliary pathways such as xylose isomerase. Subgroups 2 and 4 retain complete glycolytic and TCA cycle pathways, whereas subgroup 3, although lacking gluconeogenesis, is functionally enriched in sulfur and nitrogen metabolism, enabling niche specialization under anoxic conditions. Overall, the metabolic heterogeneity of WOR-3 appears to be closely linked to its wide environmental distribution, suggesting a potentially significant role in global biogeochemical cycling. This study not only expands the genomic repertoire of WOR-3 but also offers novel insights into its metabolic capacities and ecological roles, laying a foundation for future investigations into the evolutionary history and ecological significance of uncultivated microbial lineages. Further studies may explore the functional roles of WOR-3 across distinct ecosystems and their interactions with co-occurring microbial taxa, contributing to a more comprehensive understanding of microbial processes in natural environments.

## Data availability

The fifty-nine genomes retrieved in this study have been deposited at CNGB Sequence Archive (CNSA) with accession number CNP0007392.

## Acknowledgments

This work was supported by the National Natural Science Foundation of China (Nos. 42177178, 42377114, 42077281, 32471574 and 42307270).

---

**Fig. S1** - Phylogenetic tree for 16S rRNA gene. Distributions of sample MAGs and public database MAGs are labelled with red pentagrams and solid grey dots, bootstrap values >75 are labelled with solid grey rectangles with black borders.

**Fig. S2** - Phylogenetic tree constructed based on RnfC subunits from Anaerobes as well as Aerobes and facultative anaerobes.

**Table S1** Genomic characteristics of WOR-3 MAGs.

**Table S2** WOR-3 16S rRNA sequences traceability information.

**Table S3** Metabolic pathway gene annotations in WOR-3 MAGs.

**Table S4** Functional annotation and taxonomic classification summary of WOR-3 MAGs.

